# The BAP1 deubiquitinase complex is a general transcriptional co-activator

**DOI:** 10.1101/244152

**Authors:** Antoine Campagne, Dina Zielinski, Audrey Michaud, Stéphanie Le Corre, Florent Dingli, Hong Chen, Ivaylo Vassilev, Ming-Kang Lee, Nicolas Servant, Damarys Loew, Eric Pasmant, Sophie Postel-Vinay, Michel Wassef, Raphaël Margueron

## Abstract

In *Drosophila*, a complex consisting of Calypso and ASX catalyzes H2A deubiquitination and has been reported to act as part of the Polycomb machinery in transcriptional silencing. The mammalian homologs of these proteins (BAP1 and ASXL1/2/3, respectively), are frequently mutated in various cancer types, yet their precise functions remain unclear. Using an integrative approach based on isogenic cell lines generated with CRISPR/Cas9, we uncover an unanticipated role for BAP1 in gene activation. This function requires the assembly of an enzymatically active BAPl-associated core complex (BAP1.com) containing one of the redundant ASXL proteins. We investigated the mechanism underlying BAP1.com-mediated transcriptional regulation and show that it functions neither in synergy nor by antagonism with the Polycomb machinery. Instead, our results provide compelling evidence that BAP1.com acts as a general transcriptional co-activator.

## INTRODUCTION

BAP1 (BRCA1 Associated Protein 1) was initially characterized as a nuclear deubiquitinase regulating the function of BRCA1 ^1^. Subsequent work suggested that BAP1 does in fact interact with BARD1 (BRCA1-associated RING domain 1) and regulates its ubiquitination ^2^. Since then, a variety of proteins have been reported to interact with BAP1, including transcription factors (YY1, FOXK1/2), chromatin binders and modifiers (ASXL1/2/3, KDM1B, OGT1), the cell cycle regulator HCFC1 and DNA repair proteins (MBD5/6) ^3–9^. BAP1 enzymatic activity has been shown to regulate the ubiquitination of various proteins including gamma-tubulin ^10^, INO80 ^11^ or BRCA1 ^1^. Accordingly, BAP1 participates in diverse cellular processes, such as transcriptional regulation and the DNA damage response. However, the precise function of BAP1 in transcriptional regulation remains elusive; for instance, some studies have reported that BAP1 acts as a transcriptional activator while others have suggested that it is required for gene silencing ^3,4,12^.

In parallel to its characterization in mammals, studies in *Drosophila* identified the BAP1 ortholog Calypso as a novel Polycomb protein ^13,14^. The Polycomb Group (PcG) of proteins is essential for the maintenance of gene repression, most prominently at developmentally regulated genes. Consequently, altering PcG function affects key cellular processes such as cell fate determination, cell proliferation, and genomic imprinting ^15^. Two Polycomb complexes have been well characterized so far: the Polycomb Repressive Complex 1 (PRC1) and PRC2. PRC2 catalyzes di- and trimethylation of histone H3 on lysine 27 (H3K27me2/3), while PRC1 acts through chromatin compaction and monoubiquitination of histone H2A on lysine 119 (H2AK119ub1) ^16,17^. Conserved from *Drosophila* to mammals, the activity of these complexes is necessary for maintaining transcriptional silencing of their target genes. The importance of H2AK119ub1 in Polycomb silencing has recently been called into question in *Drosophila* ^18^ as well as in mouse models ^19^. Nonetheless, recent studies suggest that this mark participates in stabilizing PRC2 binding to chromatin ^20,21^.

*Drosophila* Calypso was found to partner with the Polycomb protein Additional Sex Combs (ASX) into a novel Polycomb complex termed Polycomb Repressive DeUBiquitinase (PR-DUB) complex. PR-DUB was shown to catalyze deubiquitination of H2AK119ub1, an activity opposite to that of PRC1 ^22^ The interaction between BAP1 and homologs of ASX (ASXL1/2/3 proteins) is conserved in mammals, as is the H2AK119 DUB activity of BAP1 ^13,23^. However, how the antagonistic activities of Calypso/BAP1 and PRC1 converge to maintain transcriptional silencing remains enigmatic. In addition, the link between PR-DUB and the Polycomb machinery is still controversial. Some studies have reported that ASXL1 interacts with PRC2 and is required for its recruitment ^24–28^ while others have suggested an antagonism between BAP1 and PRC2 ^29,30^. Altogether, a clear picture of the function of BAP1 and ASXL proteins is still lacking. Understanding the function of PR-DUB is all the more important in view of the tumor-suppressive functions of BAP1 in several cancer types including uveal melanoma, mesothelioma, and clear-cell renal cell carcinoma and of ASXL proteins in hematologic malignancies ^2,31^.

In this study, we used biochemical, genome-editing, and genome-wide approaches to address the function of BAP1 in transcriptional regulation and its relationship to the Polycomb machinery. We show that the ASXLs are mandatory partners of BAP1 and that they are required for its stability and its enzymatic activity. At the functional level, the complex formed around BAP1 (BAP1.com) is required for efficient transcription of many developmental genes, a function that depends on the deubiquitinase (DUB) activity. Accordingly, BAP1 appears to be largely dispensable for maintaining silencing of Polycomb-target genes and in fact opposes PRC2-mediated silencing at a number of genes. The majority of BAP1-regulated genes, however, are not under regulation by PRC2, suggesting that the function of BAP1 in promoting transcription does not reflect an antagonism with Polycomb proteins. Indeed, we show that BAP1 is required upon transcriptional stimulation as observed upon retinoic acid treatment, a function that is shared with the CREBBP and SMARCB1 transcriptional co-activators. A general role in regulating gene expression is supported by the conspicuous co-localization between BAP1.com and the RNA polymerase 2. Altogether, our integrative analysis uncovers an essential function for BAP1.com as a transcriptional co-activator.

## RESULTS

### BAP1 inactivation alters the expression of developmentally regulated genes

In order to comprehend the role of BAP1 in transcriptional regulation, we generated knockouts for *BAP1, ASXL1, ASXL2*, and *EZH2*, encoding the main catalytic subunit of PRC2, in human HAP1 cells, a model cell line that is nearly haploid and thus particularly amenable to genome editing ^32^. Knockouts (KO) result from the insertion of a STOP cassette that interrupts transcription and translation of the target gene ^33^ and were validated by RT-qPCR (Figure 1A). Notably, in contrast to many cell lines where the knockdown of BAP1 severely compromises proliferation (Figure S1A), we found that BAP1 is dispensable for proliferation in HAP1 cells, thus providing a suitable system for studying its mechanism of action (Figure 1B). Cellular fractionation confirmed the tight association of BAP1 with chromatin, as shown by its enrichment both in the soluble and insoluble fractions (Figure S1B), which prompted us to further investigate its chromatin-modifying activity. Western blot analysis of various histone marks showed that BAP1 loss is associated with an approximate 2-fold increase in total H2A ubiquitination levels, as well as a parallel increase in H2A.Z ubiquitination, with no effect on H2B ubiquitination (Figure 1C, top panel). *ASXL1* and *ASXL2* KO led to a milder increase in H2A/H2A.Z ubiquitination. We did not find any global effect of knocking out *BAP1, ASXL1*, or *ASXL2* on other histone marks such as H3K4me2 and H3K27me3 or on DNA modifications (Figure 1C, bottom panel).

**Figure 1:**
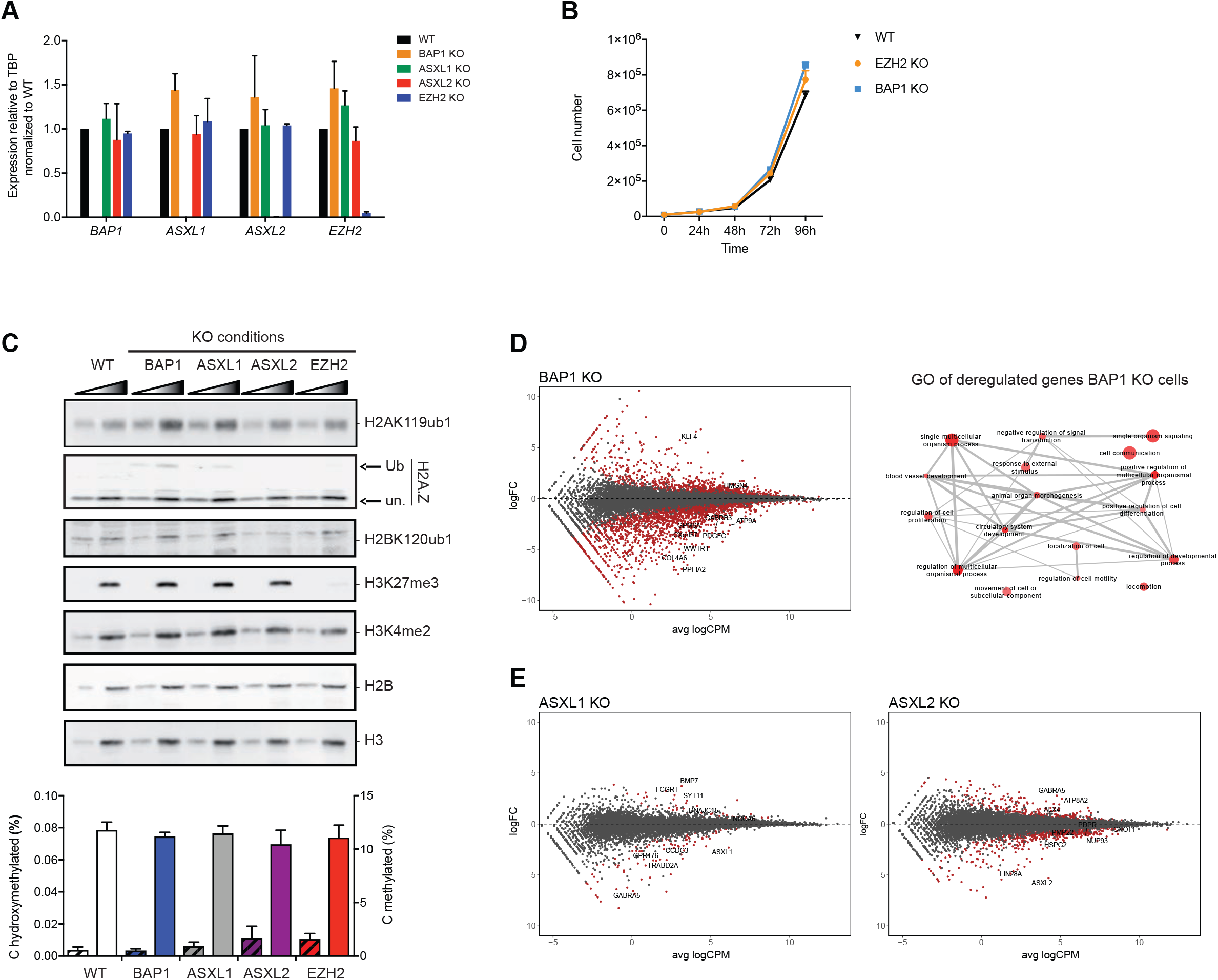
Functional consequences of loss of BAP1, ASXL1 or ASXL2 on chromatin and gene expression. **A**) RT-qPCR analysis of *BAP1, ASXL1, ASXL2* and *EZH2* expression in the different KO conditions indicated on top. Mean ± SD. n = 2. **B**) Proliferation curve of wild-type, *EZH2* KO and *BAP1* KO cells. Mean ± SD. n = 3**. C)** Top, western blot analysis of acid extracted histones with antibodies directed against various histone modifications (as indicated on the right) in the different cell lines indicated on top. A two-point titration (1:2.5 ratio) is shown for each condition. Bottom, analysis of cytosine methylation (solid bars) or hydroxymethylation (dashes) in the conditions indicated at the bottom. Mean ± SD. n = 3**. D)** Left panel: scatterplot showing log2 fold-change (logFC) expression between wild-type and *BAP1* KO cells versus average log2 counts per million (logCPM). Differentially expressed genes (DEGs) in *BAP1* KO cells are highlighted in red. Right panel: Representation of the non-redundant most enriched GO terms within the DEGs in *BAP1* KO cells**. E)** Scatterplots as in D, showing gene expression changes in *ASXL1* and *ASXL2* KO cells.

We then performed RNA-seq to analyze the transcriptome of wild-type and KO cell lines. The inactivation of BAP1 leads to dramatic changes in gene expression (1893 differentially expressed - DE genes, FDR < 0.05 and absolute log2 fold-change > 1; Figure 1D, left panel). Strikingly, a majority of the affected genes were downregulated in *BAP1* KO cells (n=1370), suggesting an involvement in gene activation rather than silencing. Gene ontology (GO) analysis revealed enrichment for a variety of biological processes ranging from broad terms such as cell communication, signaling, or regulation of proliferation to more specific terms such as blood vessel development (Figure 1D, right panel). Importantly, several terms related to development are also found enriched, which is consistent with the reported function of the BAP1 ortholog in *Drosophila* ^13^. Inactivation of either ASXL protein also leads to preferential downregulation of genes (85 downregulated genes out of 112 DE genes in *ASXL1* KO cells and 262 downregulated genes out of 406 DE genes in *ASXL2* KO cells; Figure 1E, also see Figure S1C for heatmaps of the top 100 DE genes in each KO condition) but has a much milder effect on gene expression than observed for *BAP1* KO. In addition, loss of ASXL1 or ASXL2 does not affect the localization of BAP1 at chromatin (Figure S1B). This result suggests either that BAP1 can function independently of its interaction with ASXL proteins or, alternatively, that the ASXLs are largely redundant.

### The ASXL/BAP1 core complex is conserved in mammals

To better understand the function of BAP1, we sought to determine whether it is part of a stable complex and, if it is, with which partners. The purification of Calypso from *Drosophila* embryos revealed that it forms a heterodimeric complex with ASX ^13^. While the interaction between BAP1 and ASXL proteins is conserved ^13,23,34^, additional partners were reported in the mammalian complex ^3,4,9^. We overexpressed FLAG-tagged versions of BAP1, ASXL1, or ASXL2 in HeLa cells (Figure S2A), followed by immunoprecipitation and mass spectrometry. The results confirmed previous reports, in particular the identification of ASXL1/2, FOXK1/2, HCFC1, and KDM1B as partners of BAP1 (Figure 2A). The ASXL1 and ASXL2 interactomes are similar with the exception of KDM1B, which is specifically pulled down by ASXL2. Of note, similar results were obtained for the BAP1 interactome in Uveal Melanoma MP41 cells (Figure S2B), suggesting that BAP1.com composition is not cell-type-dependent.

**Figure 2:**
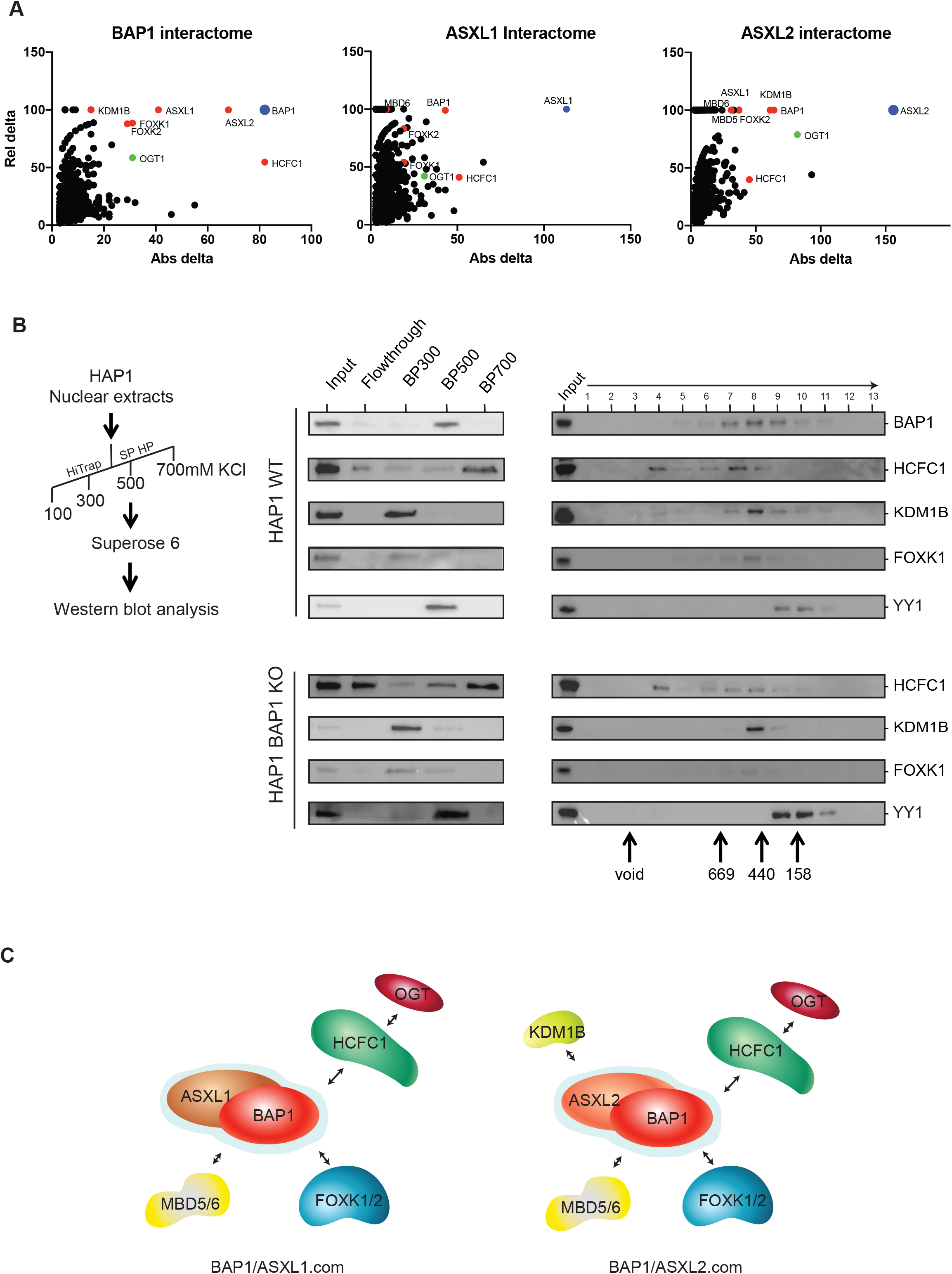
Bap1.com core complex and associated factors. **A**) Mass spectrometry analysis of HeLa cells overexpressing Flag-tagged versions of BAP1, ASXL1 and ASXL2. Graphs represent proteins relative to their absolute and relative delta compared to mass spectrometry analysis of empty vector expressing cells. Absolute delta is the absolute difference between distinct peptides identified in sample and control; relative delta is the ratio of absolute delta versus the sum of distinct peptides identified in sample and control**. B)** Elution patterns of HAP1 WT and HAP1 *BAP1-KO* nuclear extracts following the purification scheme indicated in the left panel and monitored by western blot with the indicated antibodies. Middle pattern is a representative elution pattern (step elution with increased salt concentration) on a cation exchange column (SP-HP, GE). Right panel is a representative elution pattern on a size-exclusion column (Superose 6, PC3.2/30, GE)**. C)** Schematic of BAP1.com core complex and associated factors depending on the ASXL paralog present.

We then sought to determine whether BAP1 partners are all present in a single complex or if BAP1 is engaged in distinct protein complexes. To this end, we analyzed the elution pattern of BAP1 partners by ion exchange chromatography followed by size-exclusion chromatography (SEC) (Figure 2B). The first purification step (ion exchange) revealed that the majority of HCFC1, FOXK1, and KDM1B elute independently of BAP1, which is almost exclusively found in the 500 mM salt fraction, suggesting that only a portion of each of these proteins is engaged in a complex with BAP1. In the second purification step (size-exclusion), we observed that BAP1 elutes with an apparent molecular weight of approximately 500kDa, overlapping with HCFC1, FOXK1, and KDM1B. In contrast, YY1 elutes later and only partially overlaps with BAP1. To know whether the co-elution between BAP1 and its cofactors reflect the assembly of these proteins into a complex, we repeated the experiment with a nuclear extract from BAP1 KO cells. The elution pattern from cation exchange did not reveal any major changes; we therefore continued with SEC. The elution patterns of HCFC1, KDM1B, and FOXK1 remain unchanged indicating that the co-elutions observed with BAP1 in the wild-type extract do not reflect the formation of a stable complex (Figure 2B). Considering the previously established tight interaction between BAP1 and the ASXLs ^13,23,34^, our results suggest that BAP1 and ASXL proteins form a core complex (BAP1.com) which engages in transient interactions with additional partners such as FOXK1/2, HCFC1 or KDM1B (Figure 2C). This model predicts that immunoprecipitation of any one of these transient partners would consistently retrieve the core complex but not necessarily other transient partners. Indeed, KDM1B immunoprecipitation from HeLa cells pulls down ASXL2 and BAP1 (as well as NSD3, which is part of a distinct complex with KDM1B) but none of the other transient partners (Figure S2C).

### ASXLs are required for BAP1 function

Given the role of ASXL1 and ASXL2 in driving BAP1-associated complex composition but the modest effect of their individual deletion on transcription, we wished to investigate their potential redundancy in BAP1-mediated H2A deubiquitination and transcriptional regulation.

To determine whether both proteins also participate in BAP1 function in cells, we generated a double *ASXL1/ASXL2* knockout. Of note, *ASXL3* is not expressed in wild-type HAP1 cells nor in *ASXL1/2* double knockout cells (Figure S3A). We first evaluated the effect of this double knockout on chromatin regulation and observed a robust increase in H2A ubiquitination levels similar to those observed in the *BAP1* knockout (Figure 3A). This result is consistent with the fact that interaction with the ASXLs is required for BAP1 enzymatic activity (Figure S3B). Notably, loss of ASXL1 and ASXL2 also led to a dramatic reduction in BAP1 protein levels, indicating that the ASXLs are not only necessary for the enzymatic activity of BAP1 but also for its stability *in vivo* (Figure 3A). Consistent with this effect on BAP1 protein accumulation, transcriptome analysis of *ASXL1/2* dKO cells revealed a major impact on gene expression (Figure 3B). Similarly to *BAP1* KO, most differentially expressed genes were downregulated (1073 downregulated genes out of 1576 DE genes, also see Figure S3C for heatmaps of the top 100 DE genes) and there was a large overlap between genes downregulated in *BAP1* KO and *ASXL1/2* dKO cells (Figure 3C). In comparison, the overlap between upregulated genes was much less pronounced (Figure 3D), supporting the idea that the main role of BAP1.com is to promote transcription.

**Figure 3:**
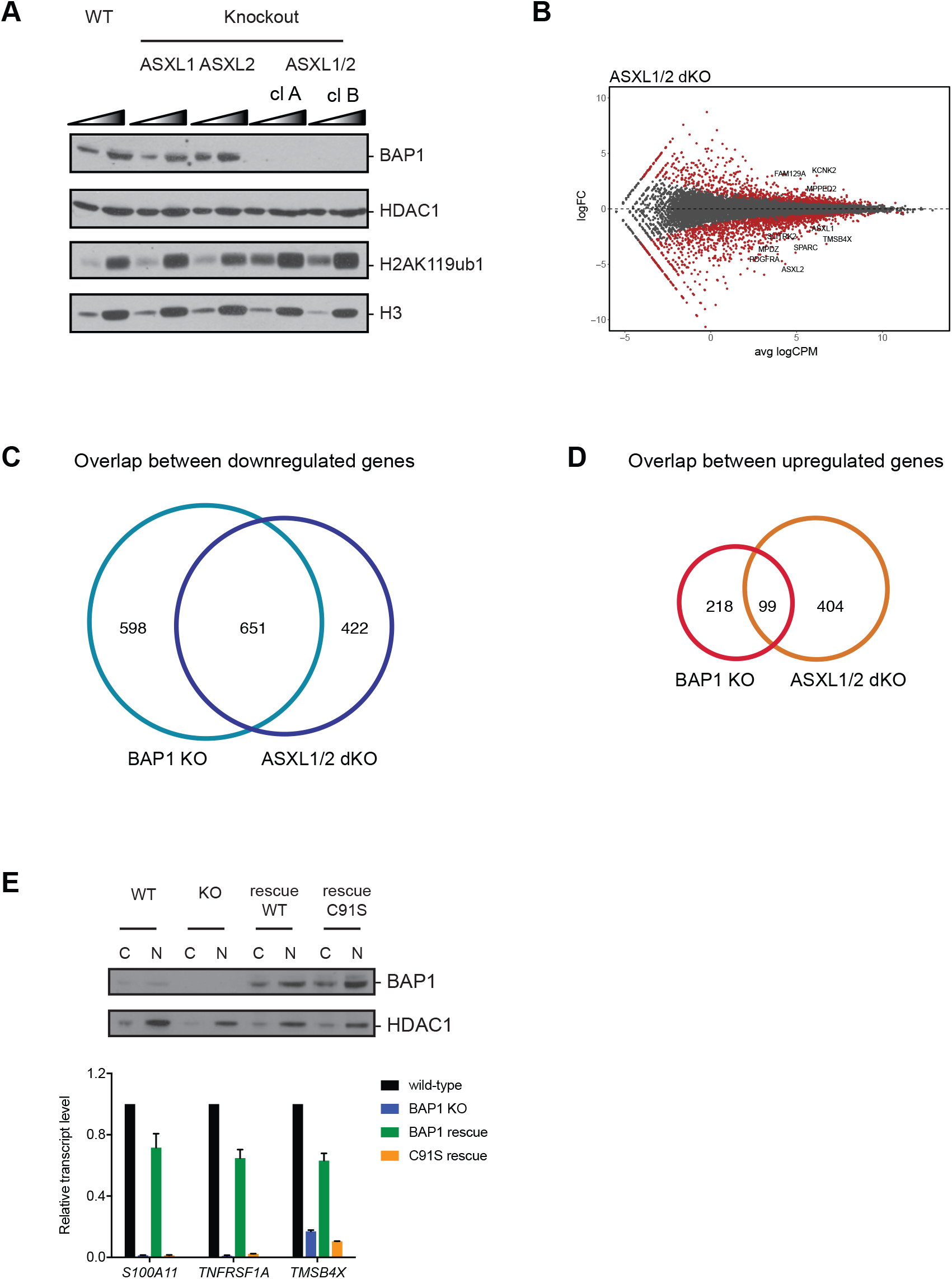
requirement of ASXL1 and ASXL2 and of the DUB activity for BAP1 function. **A**) Western blot analysis of BAP1 and H2AK119ub1 in the different KO conditions indicated above. 2 independent clones of *ASXL1/2* dKO cells are shown. HDAC1 and H3 serve as nuclear and histone protein loading control respectively. A two-point titration (1:2.5 ratio) is shown for each condition**. B)** Scatterplot showing log2 fold-change (logFC) expression between wild-type and *ASXL1/2* dKO cells as a function of average log2 counts per million (logCPM). Differentially expressed genes in *ASXL1/2* dKO cells are highlighted in red. **C, D**) Venn diagram showing the overlap between genes downregulated (**C**) or upregulated (D) in *BAP1* KO and *ASXL1* /2 dKO cells**. E)** Reintroduction of wild-type or catalytically-dead (C91S) BAP1 in *BAP1* KO cells. Top panel: western blot analysis of BAP1 expression in cytoplasmic (C) or nuclear (N) fractions. HDAC1 is used as a loading control. Bottom panel: RT-qPCR analysis of three BAP1-regulated genes in wild-type, *BAP1* KO and the two rescue conditions (Mean ± SD, n = 3).

We then wondered if the function of BAP1 in regulating gene transcription requires its deubiquitinase activity. For this purpose, we performed rescue experiments in *BAP1* KO cells, reintroducing either wild-type BAP1 or a catalytically dead version (BAP1 C91S). Both versions of BAP1 were re-expressed at a similar level, slightly higher than the original endogenous level (Figure 3E, upper panel). Focusing on a selection of genes whose expression is dramatically reduced in the absence of BAP1, re-expression of wild-type BAP1 protein restores up to 75% of the wild-type levels of the transcripts while the C91S mutant is unable to rescue transcription of the tested genes (Figure 3E, lower panel). Altogether, these results demonstrate that BAP1 associates with the ASXL proteins to promote transcription in a manner that depends on its enzymatic activity.

### BAP1 does not participate in Polycomb-mediated silencing

The data presented above suggest that the main role of BAP1 is to positively regulate transcription. While this is in agreement with several previously published studies, it contrasts with a number of reports suggesting that BAP1 and ASXL proteins participate in Polycomb-mediated silencing ^24–28^. To formally investigate the interplay between BAP1 and Polycomb proteins, we analyzed the consequences of BAP1 loss in conjunction with loss of RING1B and EZH2, key members of PRC1 and PRC2, respectively (Figure 4A).

**Figure 4:**
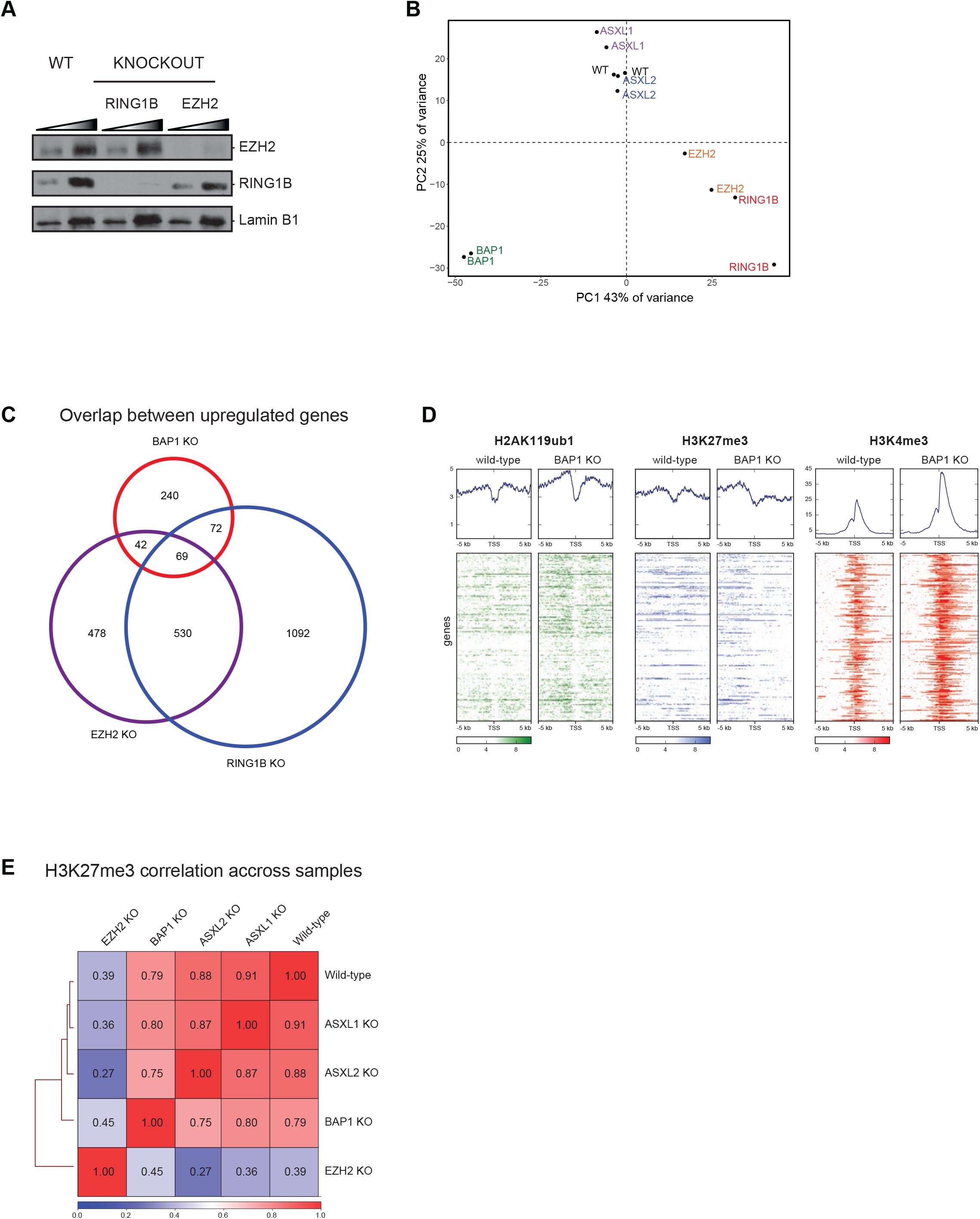
BAP1.com is dispensable for Polycomb-mediated silencing. **A**) Western blot analysis of EZH2 and RING1B in wild-type (WT), *RING1B* KO, and *EZH2* KO cells. Lamin B1 is used as a loading control. A two-point titration (1:2 ratio) is shown for each condition**. B)** PCA analysis of WT, *RING1B, EZH2, ASXL1, ASXL2*, and *BAP1* KO transcriptome**. C)** Venn diagram showing the overlap between genes upregulated in *BAP1, EZH2* or *RING1B* KO cells**. D)** Heatmaps showing H2AK119ub1, H3K27me3, and H3K4me3 distribution in a -5/+5 kb window around the transcription start site (TSS) of genes upregulated in *BAP1* KO cells in wild-type and *BAP1* KO cells. Corresponding average profiles are plotted on top of each heatmap**. E)** Correlation heatmap of H3K27me3 distribution between the different KO conditions indicated.

As expected, the main impact of inactivating either *RING1B* or *EZH2* is the transcriptional upregulation of a large set of genes (Figure S4A and S4B). The GO terms of the differentially expressed genes (either up- or downregulated) in EZH2 KO cells partially overlap the categories observed upon BAP1 KO such as terms related to signaling or development (Figure S4C to Figure 1D, right panel). The terms associated with genes differentially expressed in the absence of RING1B are broader, which might reflect a less developmental specific function of PRC1 (Figure S4D). To further compare transcriptional changes among all KO conditions, we performed a principal component analysis (PCA, Figure 4B). Along the main dimension of the PCA, *BAP1* KO and Polycomb KO cells clearly vary in opposite directions, again suggestive of distinct gene regulatory functions. Since the Polycomb machinery is involved in gene silencing, we investigated the overlap between genes upregulated upon KO of *EZH2* or *RING1B* and KO of *BAP1*. As shown in Figure 4C, only a minority of the genes regulated by PRC1 and/or PRC2 become upregulated upon loss of BAP1. To determine whether this limited overlap nonetheless reflects a synergistic action of BAP1 with the Polycomb machinery, we investigated chromatin changes occurring at genes upregulated in *BAP1* KO. If BAP1 functions together with the Polycomb machinery to maintain transcriptional silencing at a subset of Polycomb target genes, loss of BAP1 is expected to result in a decrease in the Polycomb-mediated chromatin signature. However, chromatin-immunoprecipitation followed by deep sequencing (ChIP-seq) did not reveal a decrease in Polycomb histone marks H2AK119ub1 or H3K27me3 at these genes in *BAP1* KO cells (Figure 4D). In fact, H2AK119ub1 increased, possibly reflecting the more global increase of the mark caused by loss of BAP1. As expected, transcriptional upregulation corresponded with marked increase in H3K4me3, a histone mark deposited preferentially near the 5’ ends of transcriptionally active genes. These results suggest that transcriptional upregulation occurring upon BAP1 loss is not caused by impaired Polycomb-mediated silencing. Instead we believe that these gene expression changes are secondary effects of widespread transcriptional downregulation.

Finally, since previous studies have reported a crucial role for the ASXLs in H3K27me3 deposition ^24–28^, we further analyzed the genomic distribution of H3K27me3 in wild-type, *BAP1, ASXL1, ASXL2*, and *EZH2* KO conditions. As shown in Figure 4E, there is a good genome-wide correlation in the localization of the mark between wild-type, *BAP1, ASXL1* and *ASXL2* KO cells, suggesting that loss of the ASXL proteins does not globally affect H3K27me3 distribution (see also Figure S4E for an example snapshot). This analysis, together with the lack of change in H3K27me3 abundance in *BAP1, ASXL1*, or *ASXL2* KO cells (Figure 2C), rules out an essential role for BAP1 and ASXL proteins in PRC2 function, challenging the notion that BAP1 critically participates in Polycomb-mediated silencing.

### Polycomb-antagonistic and -independent role of BAP1 in promoting transcription

As shown above, the major impact of loss of BAP1 is the downregulation of gene expression. A straightforward explanation of this effect on gene transcription lies in BAP1 deubiquitinase activity of the PRC1-deposited histone mark, H2AK119ub1. Although H2AK119ub1 has been suggested to be dispensable for gene silencing, it has also been reported to stimulate PRC2 recruitment and to locally promote H3K27me3 deposition ^20,21,35^.

To investigate whether BAP1 inactivation indeed results in a gain in Polycomb function, we analyzed genes that are downregulated in the absence of BAP1 (Figure 5A). As expected, H3K4me3 decreased concomitantly with transcriptional downregulation. In contrast, both H2AK119ub1 and H3K27me3 significantly increased. In principle, increased levels of these repressive marks could be a direct consequence of the loss of BAP1 deubiquitinase activity or a secondary event due to transcriptional downregulation ^36^. In the first case, the Polycomb machinery is expected to contribute to transcriptional silencing, whereas in the second case it is unlikely to be required. To discern between these two possibilities, we genetically inactivated *EZH2* in *BAP1* KO cells (Figure 5B) to determine whether EZH2 deletion can restore the expression of genes that have been downregulated as a consequence of BAP1 loss. Of note, *EZH2* deletion in cells already KO for *BAP1* did not impair proliferation (Figure 5C), consistent with recent evidence challenging the reported synthetic lethal relationship between EZH2 inhibition and BAP1 inactivation ^29,37^. More importantly, of the 913 genes downregulated upon *BAP1* KO, a large majority (741 genes) remain silent in the *BAP1/EZH2* double KO (Figure 5D, top heatmap) while the expression of a minority (172 genes) is increased in the double KO compared to BAP1 single KO cells (Figure 5D, bottom heatmap). This set of genes is also upregulated upon deletion of *EZH2* in a *BAP1* wild-type context, indicative of a balanced antagonistic regulation by BAP1 and EZH2 (Figure 5D and 5E). Moreover, all genes downregulated in the absence of BAP1 (i.e. whether or not they are sensitive to loss of EZH2) gain H2AK119ub1 and H3K27me3 upon *BAP1* knockout (Figure 5F, see also Figure 5G and Figure S5 for specific examples), suggesting that this gain in repressive marks is a consequence rather than a cause of transcriptional downregulation. Thus, although BAP1 and Polycomb proteins act in an opposite fashion at a number of genes, BAP1 promotes gene expression in a way that is largely independent of an antagonism with the Polycomb machinery.

**Figure 5:**
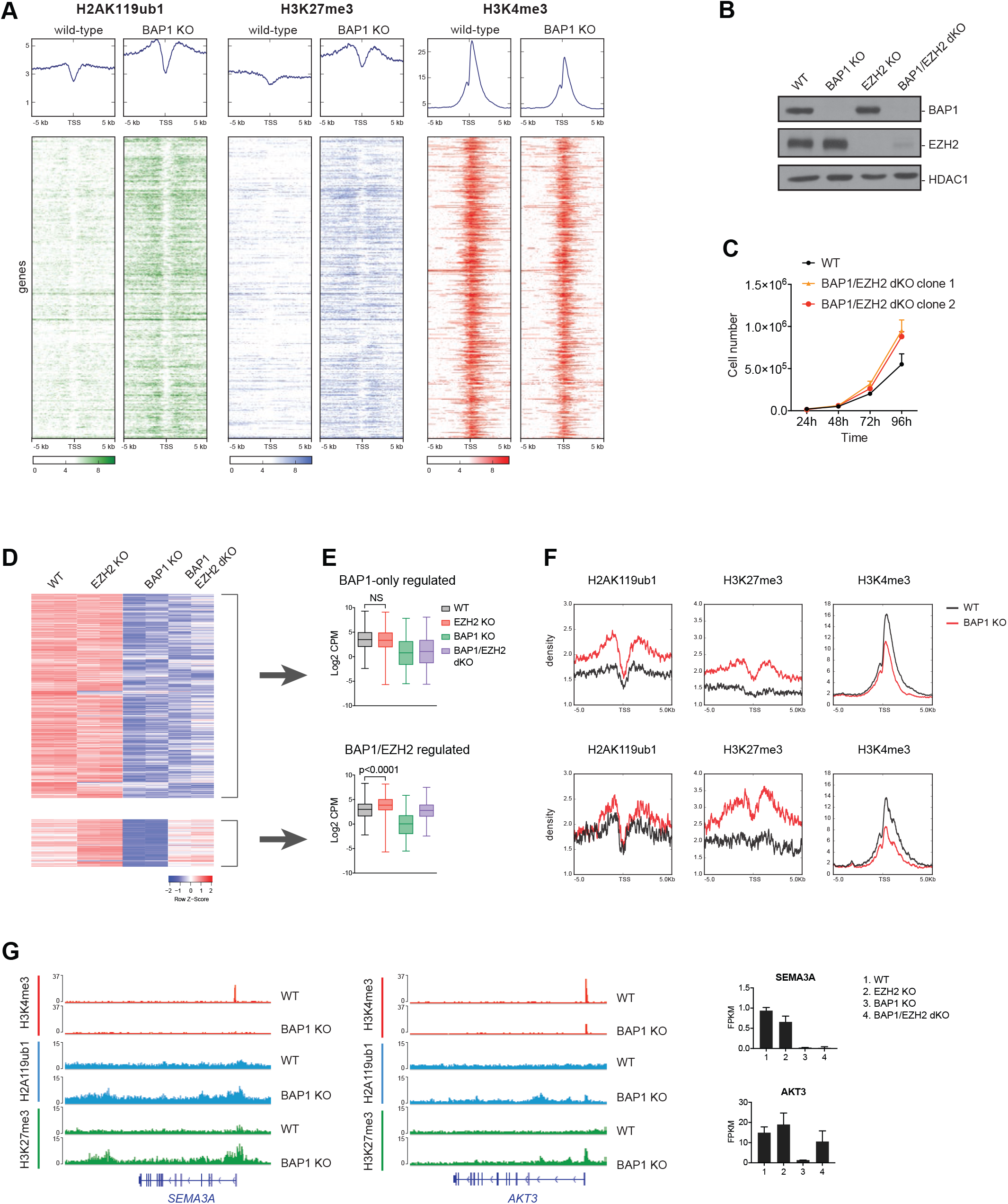
BAP1.com promotes transcription independently of an antagonism with Polycomb proteins. **A**) Heatmaps showing H2AK119ub1, H3K27me3 and H3K4me3 distribution in a -5/+5 kb window around the transcription start site (TSS) of genes downregulated in *BAP1* KO cells in wild-type and BAP1 KO cells. Corresponding average profiles are plotted on top of each heatmap**. B)** Western blot analysis of BAP1 and EZH2 in single and double *BAP1/EZH2* KO cells. HDAC1 is used as a loading control**. C)** Proliferation curve of wild-type cells and two independent clones of *BAP1/EZH2* dKO cells. Mean ± SD. n = 3**. D)** Heatmap of gene expression (Z-scores) in the different genotypes. Top: BAP1-only-regulated genes, and bottom: BAP1- and EZH2-regulated genes**. E)** Box-plots of log2 CPM expression values of BAP1-only-regulated genes (top) and BAP1/EZH2-regulated genes (bottom) in the different conditions as indicated. Result of the Mann–Whitney test on the *EZH2* KO versus wild-type comparison is indicated**. F)** Plot showing average enrichment of H2AK119ub1, H3K27me3, and H3K4me3 in a -5/+5 kb around the TSS for BAP1-only-regulated genes (top) and BAP1/EZH2-regulated genes (bottom)**. G)** Example snapshots of H2AK119ub1, H3K27me3, and H3K4me3 enrichment in WT and *BAP1* KO cells at a BAP1-only-regulated gene and a BAP1/EZH2-regulated gene (middle). Expression of the corresponding genes across WT, *EZH2, BAP1*, and *BAP1/EZH2* KO conditions as detected in corresponding RNA-seq data is shown on the right.

### Functional overlap between BAP1.com and general co-activators

Having analyzed the impact of loss of BAP1.com on steady state gene transcription, we next sought to examine its role in transcriptional activation in response to a transcriptional stimulus. Considering previous reports suggesting that BAP1 may modulate nuclear receptor mediated gene regulation ^38,39^, we investigated whether deletion of *BAP1* would affect response to retinoic acid (RA) treatment. We initially performed a time-course analysis of two of the best-characterized direct RA transcriptional targets: *RARβ* and *CYP26A1* (Figure 6A). In contrast to the above-mentioned reports that suggest a repressive role for BAP1.com at RA target genes, we observed that activation of both *RARβ* and *CYP26A1* was severely compromised in *BAP1* KO cells. Of note, loss of BAP1 did not affect the protein levels of RARα, a major RA-binding nuclear receptor (Figure S6A). To determine whether these results reflect a general role for BAP1 in the transcriptional response to RA, we analyzed the transcriptome of WT and *BAP1* KO cells in response to RA treatment (after 24 h). Nearly twice as many genes were significantly activated upon RA treatment in WT than in *BAP1* KO cells (n=88 versus n=47, FDR < 0.05 and absolute log2 fold-change > 1, figure S6B). Furthermore, analyzing the entire set of genes activated in either condition, we found that the response to RA is significantly attenuated in *BAP1* KO cells (Figure 6B), demonstrating that BAP1 is required for optimal RA-mediated transcriptional activation.

**Figure 6:**
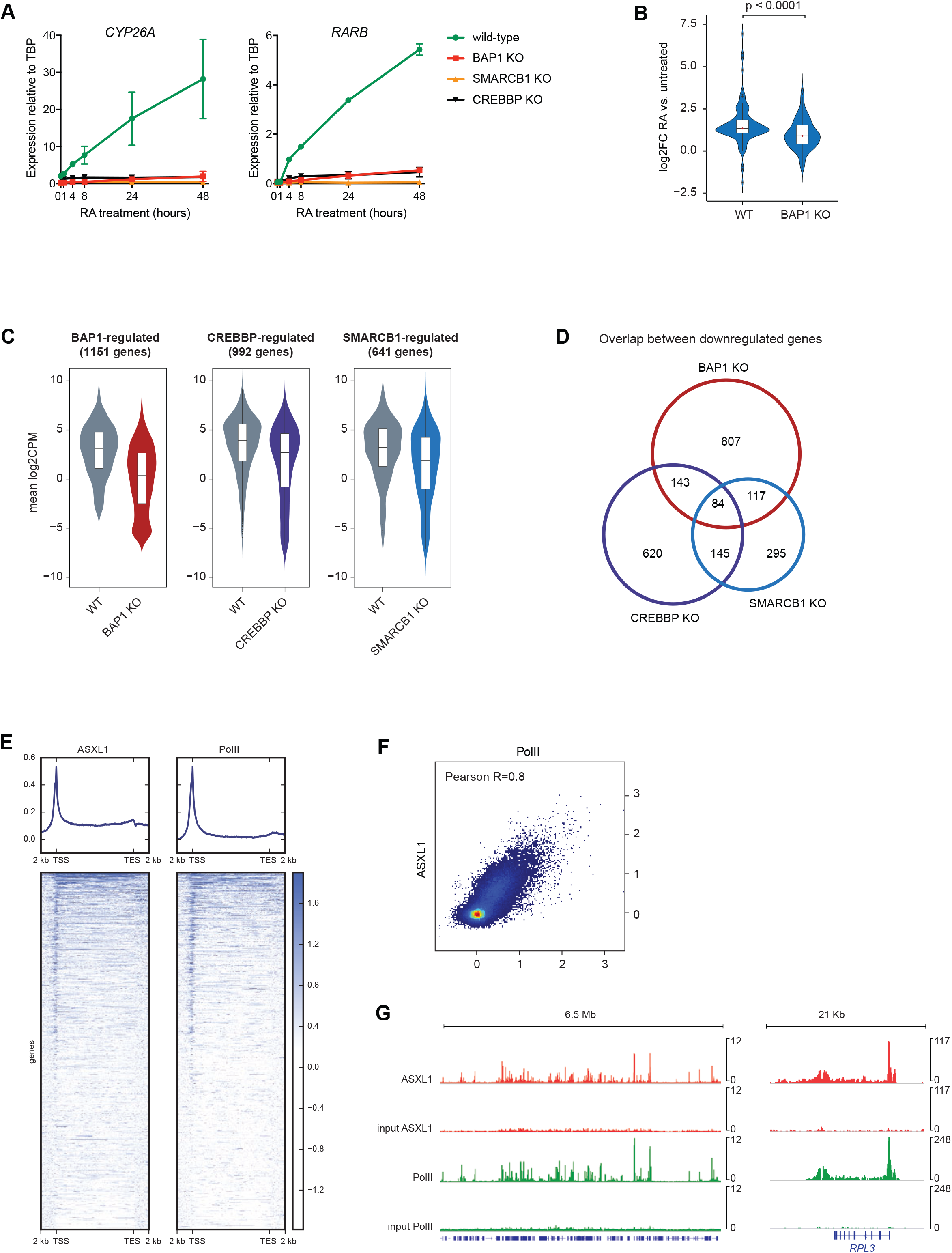
comparison of BAP1 and the CREBBP and SMARCB1 coactivator proteins. **A**) RT-qPCR analysis of *CYP26A1* and *RARB* expression following RA treatment at different time-points in *BAP1, CREBBP*, and *SMARCB1* KO cells. Mean ± SD. n = 2**. B)** Violin plots showing log2 fold-change expression of RA-responsive genes (n=114 genes, see text for details) in wild-type and BAP1 KO cells. P-value from the Mann–Whitney test is shown**. C)** Violin plots showing log2CPM expression of BAP1-CREBBP- and SMARCB1-regulated genes in wild-type (WT) or in the respective KO conditions**. D)** Venn diagram showing overlap between genes that are downregulated in *BAP1*, *CREBBP* and *SMARCB1* KO cells E) Heatmaps showing ASXL1 and RNA PolII density around the TSS and termination end site (TES) (+/- 2kb) scaled to an equivalent 10 kb. Corresponding average profiles are plotted above each heatmap**. F)** Scatterplot showing PolII versus ASXL1 enrichment around the TSS (+/- 2 kb) at all annotated genes. Pearson correlation coefficient is displayed**. G)** Snapshots of ASXL1 and RNA PolII enrichment at representative regions. The input is displayed below each corresponding ChIP-seq experiment.

This result suggests that BAP1 functions as a transcriptional co-activator that is required for efficient transcriptional induction. To further investigate this hypothesis, we compared transcriptional defects occurring in *BAP1* KO cells and knockouts of *SMARCB1* (Figure S6C), which encodes an essential member of the BAF chromatin-remodeling complex, or *CREBBP*, which encodes a histone acetyltransferase. Both KO cell lines were viable, although they appear unhealthy and have a slower proliferation rate (data not shown). As with loss of BAP1, loss of SMARCB1 and CREBBP severely impaired RA-mediated transcriptional activation of *RARβ* and *CYP26A1* (Figure 6A). To determine to what extent BAP1, CREBBP and SMARCB1 affect transcription, we compared the transcriptome of HAP1 cells mutated for each of these genes (this study and ^40^). As expected, each KO leads to the downregulation of a large set of genes (BAP1: 1151 genes, CREBBP: 992 genes and SMARCB1: 641 genes). While the mean expression in wild-type cells of each set of genes is similar, the consequence of *BAP1* KO on transcription is more pronounced (Figure 6C). We next compared the overlap between genes regulated by the 3 proteins. Although RNA-seq for each KO condition was performed in different laboratories, there was a significant overlap between genes downregulated upon loss of BAP1, CREBBP, or SMARCB1 (Figure 6D, p<0.0001 for all three comparisons). Nonetheless, the majority of downregulated genes remain specific to each respective knockout, indicating that BAP1, CREBBP, and SMARCB1 each regulate a distinct set of genes.

To determine the level of specificity of gene recruitment, we compared BAP1.com localization at chromatin to the localization of transcription machinery. We made use of previously published ChIP-seq data of either ASXL1 ^9^ and RNA-PolII (ENCODE) in HEK-293 cells, or BAP1 ^4^ and RNA-PolII (ENCODE) in mouse Bone Marrow-Derived Macrophages. Analysis of ASXL1/PolII and BAP1/PolII enrichment revealed a marked correlation between RNA-PolII and BAP1.com profiles, both in terms of enrichment intensity and profile along the gene body (Figures 6E and 6F and Figures S6D and S6E and example ChIP-seq snapshots in Figures 6G and S6F). This result strongly supports a role for BAP1.com as a general co-activator and suggests that functional differences between BAP1 and other co-activators are due to gene-specific requirements for their respective enzymatic activities during transcriptional activation, rather than gene-specific targeting. Together, these data provide compelling evidence that BAP1.com functions as a general transcriptional co-activator.

## DISCUSSION

Despite their prominent role as tumor suppressors, the biological functions of BAP1 and ASXL proteins remain poorly characterized. In this study, we performed extensive biochemical and genetic analyses in isogenic mutant cell lines to address three outstanding questions: 1) what is the role of BAP1 in transcriptional regulation? 2) what are the respective contributions of BAP1 and the ASXL to the regulation of gene expression? 3) what is the interplay between BAP1.com and Polycomb machinery?

Both biochemical and genetic evidence presented here indicate that BAP1 and ASXL proteins function together to regulate H2AK119ub1 and gene expression. Our biochemical analyses largely confirm the previously reported interactome of BAP1 ^9^ and further enable distinguishing transient interactors (FOXK1/2, MBD5/6, HCFC1, etc.) from the core complex composed of BAP1 and one ASXL paralog. With the exception of KDM1B, which appears to interact specifically with BAP1-ASXL2, the BAP1 interactome is essentially identical whether the complex forms around ASXL1 or ASXL2. This might reflect the fact that most of the interactions are mediated directly by BAP1, as suggested for HCFC1, YY1, OGT, and FOXK1/2 ^3,34^. Nonetheless, the interaction with the ASXL paralogs is required for BAP1 enzymatic activity and protein stability. The overall interchangeability between the ASXLs explains the observation that knocking out a single paralog only mildly affects gene expression. An immediate implication of this finding concerns the tumor-type-specific spectrum of BAP1 and ASXL mutations. While previous studies have argued that this non-overlapping mutation spectrum might be the result of independent and sometimes opposite functions of BAP1 and ASXL proteins ^24,29^, our findings instead suggest that, due to the redundancy among ASXL proteins, the loss of any one of them results in a much less severe disruption of BAP1.com function than loss of BAP1. Hence, we propose that the predominance of ASXL mutations in myeloid malignancies may be the result of a selective pressure aimed at only partially ablating BAP1 function, while loss of BAP1 in other malignancies would reflect a need for complete inactivation of the complex.

In contrast to the situation in *Drosophila*, our results argue against an implication of BAP1.com in the Polycomb machinery. Instead, the picture that emerges from this work is that the function of BAP1.com is to promote transcription. This property may not be surprising given that the activity of the complex is opposite to that of PRC1 in the regulation of H2A ubiquitination and that PRC1 and ASXL1 show rather distinct localization patterns ^9^. Our findings are also in line with previous studies using artificial recruitment of BAP1 to a transgene ^3^, overexpressing hyperactive forms of ASXL1 ^41^ or focusing on individual loci ^30^. We observed that a set of genes is antagonistically regulated by BAP1.com and the Polycomb machinery. However, this antagonism only occurs at a minority of BAP1-regulated genes and both regulatory activities seem to act independently of each other. Overall, BAP1.com mostly regulates the expression of genes that are not under the regulation of Polycomb proteins. As such, BAP1 does not appear to act as an anti-repressor, as described for the Trithorax and Ash1 histone methyltransferases ^42^, but rather as a general transcriptional co-activator. An obvious question arising from these results is whether BAP1 and Calypso have divergent functions or whether we can reconcile their proposed contributions to gene regulation in both species. First, it is noteworthy that there is no consensus as to whether the Calypso-Asx complex is generally involved in Polycomb-mediated silencing. Indeed, mutation of Asx in *Drosophila* leads to a complex phenotype that exhibits features of both Polycomb and Trithorax mutants ^43^, a situation that is also found in Asxl1 mutant mouse embryos ^44^. Second, only a subset of Polycomb target genes was found to be aberrantly activated upon loss of Asx or Calypso ^45^. Third, part of the difference might result from the extent to which BAP1 and Calypso regulate H2A ubiquitination. While loss of Asx leads to an approximate 10-fold increase in the total levels of H2AK118ub1 ^22^, we only observe an approximate 2-fold increase in the level of the mark upon inactivation of either of BAP1 or ASXL1/2. This suggests that in *Drosophila*, the Calypso-Asx complex might have a more critical function in restricting PRC1 activity to its normal site of action. Aberrant deposition of H2AK119ub1 and consequently of H3K27me3 could indirectly impair Polycomb-mediated transcriptional silencing. Genome-wide functional studies analyzing the global consequences of loss of Asx or Calypso in fly should help clarify these questions and address whether PR-DUB and BAP1.com have similar functions.

In addition to demonstrating that BAP1.com promotes transcription independently of an antagonism with Polycomb proteins, our analyses reveal that the response to retinoic acid treatment is similarly affected in the absence of BAP1 as upon deletion of well-known general co-activators (SMARCB1 and CREBBP). Yet, loss of each of these factors has overlapping but distinct consequences on global gene expression profiles. Considering that our analysis of previously published ChIP-seq of ASXL1 and BAP1 reveals that BAP1.com localization generally reflects transcriptional machinery genome-wide, we speculate that while present at most transcribed regions, BAP1.com impacts gene expression selectively depending on the chromatin environment. Further investigation will be necessary to decipher what determines the transcriptional response to BAP1 deletion. It will also be interesting to investigate what controls BAP1.com targeting to transcribed regions. We envision that the ASXL proteins, through their PHD finger, a domain that can potentially bind methylated lysine or arginine residues ^46^, could participate in reading post-translational marks associated with active transcription.

In conclusion, our work provides novel insight into the function of BAP1.com and paves the way for further work to elucidate the molecular mechanism linking BAP1 activity to stimulation of transcription.

### Materials and methods

#### Cell lines

HAP1 cells were kindly provided by Dr T. Brummelkamp and cultured in IMDM media supplemented with 10% FBS and 1% L-Glutamine. MP41 cells were kindly provided by Dr S. Roman-Roman and cultured in RPMI media supplemented with 10% FBS and 1% L-Glutamine (Invitrogen). HeLa-S3 cells were kindly provided by Dr S. Ait-Si-Ali. They were cultured in adherence in DMEM media supplemented with 10% FBS and 1% L-Glutamine. Non-adherent culture of HeLa cells was performed in DMEM media supplemented with 5% FBS and 1% L-Glutamine following guidelines from Nakatani and Ogryzko ^47^. SF9 cells were cultured in SF-900 II SFM medium (Invitrogen) supplemented with 5% FBS, 1% Penicillin/Streptomycin (Invitrogen) and Amphotericin B at 28° C.

#### Constitutive knockouts in HAP1 cell line

Mutations of *BAP1, ASXL1, ASXL2, EZH2*, and *SMARCB1* in HAP1 cells were performed using CRISPR/CAS9 technology as described in ^33^. The selected clones were thus used as constitutive knockouts, using the parental cell line as a control in all experiments. *RING1B* and *CREBBP* KO HAP1 cells were purchased from Horizon Discovery.

#### Stable expression in HeLa cells

For mass spectrometry analysis of Flag-tagged BAP1, ASXL1, ASXL2, and KDM1B in HeLa cells, cDNAs encoding the different proteins were first subcloned in pRev retroviral plasmid (gift from S. Ait-Si-Ali), downstream a 2xFlag-2xHA sequence and upstream an IRES sequence followed by CD25 cDNA. Retroviruses were produced by transfection of a 293 Phoenix cell line (gift from S. Ait-Si-Ali) and HeLa-S3 cells were infected by incubation with viral supernatants for 3 hours at 37°C. Infected cells were then selected by FACS sorting against CD25 expression using CD25-FITC antibody and following manufacturer’s instructions (BD Biosciences 553866). Expression was assessed by western blot analysis of nuclear extracts.

#### Rescue experiment (BAP1 WT and C91S)

Reintroduction of wild-type or enzymatically dead (C91S) BAP1was performed by infection of BAP1 KO cells with a pBABE retrovirus produced as previously described ^48^.

#### Proliferation assays

50000 cells were plated in 6 well plates in triplicates and counted every 24 hours over 4 days using a Vi-cells counter (Beckman-coulter).

#### RT-qPCR

Total RNA was isolated using Trizol-Chloroform extraction and isopropanol precipitation. cDNA was synthetized using High Capacity cDNA RT kit (4368814-Applied Biosystems) and quantitative PCR was performed with technical triplicate using SYBR green reagent (Roche) on a ViiA7 equipment (Applied Biosystems). At least three independent experiments (biological replicate) were performed for each assay and RT negative controls were always included. Primer sequences for qPCR analysis are provided in supplemental table 1.

#### RNA sequencing

Two biological independent RNA sequencing were performed for each condition. 100bp single end reads were generated for the retinoic acid analysis and 100bp paired-end reads for all other samples using the HiSeq 2500 platform. Raw reads were trimmed with cutadapt (1.12; ^49^) using the Trim Galore! (0.4.4; bioinformatics.babraham.ac.uk) wrapper (default settings) and subsequently aligned to the complete human ribosomal RNA sequence with bowtie (1.2; ^50^). Reads that did not align to rRNA were then mapped to the human reference genome (GRCh37/hg19) and gene counts generated with with STAR (2.5.2b; ^51^) using the following parameters: --quantMode GeneCounts --runThreadN 8 -- outSAMtype BAM SortedByCoordinate --runMode alignReads --outFilterMismatchNmax 6 -- outFilterMultimapNmax 20 --outSAMmultNmax 20 --outSAMprimaryFlag OneBestScore. Uniquely mapped reads are reported from the STAR log file. Total number of reads for each sample is provided in supplemental table 2.

BAM files for the CREBBP-KO and HAP1-WT samples were obtained from the Institut de Cancérologie Gustave Roussy and gene counts for these samples were generated using featureCounts (1.5.1; ^52^) with the following parameters: -C -p -s 2 -T 8 -F GTF -t exon. Raw reads were trimmed as part of bcl2fastq for Illumina adapters and aligned with RSEM (1.2.25; ^53^) and Bowtie2 (2.2.6; ^50^) to GRCh37/hg19 using default parameters. Total uniquely mapping reads were calculated using RSeQC (2.6.4 ; ^54^).

The GTF file was obtained from gencodegenes.org (comprehensive gene annotation Release 19/GRCh37.p13).

#### Differential Expression Analysis

Genes with 0 counts across all samples and CPM < 0.5 in less than 3 samples were removed. Raw library sizes were scaled using the TMM method with edgeR (3.18.1; (McCarthy, 2012 #15234)) and counts were transformed to log2-counts per million (logCPM) with limma (3.32.7). A linear model was fit to the normalized data and empirical Bayes statistics were computed for each knockout versus wildtype or rescue. Differentially expressed genes were identified from the linear fit after adjusting for multiple comparisons and filtered to include those with FDR < 0.05 and absolute logFC > 1.

Gene ontology enrichment analysis was performed with goseq (1.28.0) using the Wallenius method to calculate a probability weighting function for top differential genes as a function of a gene’s median transcript length. GO terms with FDR < 0.01 were collapsed in REVIGO.

#### CREBBP, SMARCA4, BAP1 Analysis

Raw data for SMARCA4 and WT were downloaded from GEO (GSE75515). RNA-seq analysis was performed as described previously.

#### Chromatin immuno-precipitation

ChIPs were performed as described previously ^55^. Cell confluence and amount of starting material were kept constant by plating defined number of cells two days before cross-linking.

#### ChIP sequencing

100bp single-end reads were generated using the HiSeq2500 platform. Reads were mapped to the human reference genome (GRCh37/hg19) with Bowtie2 (2.2.9) using default parameters. PCR duplicates were removed with Picard Tools MarkDuplicates (1.97; http://broadinstitute.github.io/picard): VAL IDATION_STRIN GENCY= SILENT REMOVE_DUPLICATES=true. Total uniquely mapping reads were calculated using RSeQC bam_stat.py (2.6.4). bigWig files were generated with deepTools bamCoverage (2.4.1). Read counts were normalized using a scale factor of 1000000/total reads per sample and artifact regions were excluded. The merged consensus blacklist was obtained from the Kundaje lab at Stanford University (USA).

Scores 5kb upstream and downstream of TSSs were computed from normalized bigWig files with deepTools computeMatrix (2.4.1) using reference-point mode. TSS plots were generated with deepTools plotHeatmap (2.4.1).

#### PolII, BAP1 and ASXL1 ChIP-seq analysis

Raw data were downloaded from GEO (GSE40723, GSE36027, GSE51673, and GSE31477). FASTQ files were merged for samples with multiple runs and mapping was performed as previously described to hg19 or mm10. bigWig files were generated with deepTools bamCoverage (2.4.1). Artifact regions were excluded and read counts were normalized to log2(sample/input).

The merged consensus blacklists for hg19 and mm10 were obtained from the Kundaje lab at Stanford University (USA): ftp://encodeftp.cse.ucsc.edu/users/akundaje/rawdata/blacklists/

Average values per TSS +/- 2kb were computed using deepTools multiBigwigSummary and correlation plots for these regions were generated with deepTools plotCorrelation (--corMethod pearson –removeOutliers –skipZeros).

#### Antibodies

BAP1 (C-4; sc-28383), FOXK1 (G-4; sc-373810), YY1 (sc-7341) and RAR-alpha (sc-551X) were purchased from Santa-Cruz; FLAG (M2; F1804) was purchased from Sigma; HCFC1 (A301–400A) and RNF2 (A302–869A) were purchased from Bethyl Laboratories; Lamin B1 (ab16048); H2AK119ub (D27C4; 8240S), H3K27me3 (C36B11, 9733S), H3K4me3 (C42D8, 9751) were purchased from Cell Signaling; H2B (5HH2–2A8; 61037); H2BK120Ub (C56; 39623), H3 (C-terminal; 39163), KDM1B (61457) were purchased from Active Motif; H3K4me2 (MCA-MAB10003–100-Ex) was purchased from Cosmo Bio; alpha-Tubulin (1F4E3; A01410) was purchased from Genscript.

#### Histone extraction

Cells were lysed in a hypotonic lysis buffer (10mM Tris pH6.8, 50mM Na2SO4, 1%Triton X-100, 10mM MgCl2, 8.6% sucrose and protease inhibitors) and carefully homogenized using a Dounce A homogenizer. After a 6000g 10min 4° C centrifugation step, the pellet was washed with 10mM Tris pH7.5, 13mM EDTA and resuspended in ice cold water. Protein precipitation was performed by addition of sulfuric acid 0.4N final concentration and 1hr incubation on ice. The samples were centrifugated 10min 20000g 4°C and the histone-containing supernatant collected and neutralized by addition of 0,5 volume of 1.5M Tris pH8.8. Quantification was performed by Bradford assay and confirmed by SDS-PAGE stained with Coomassie.

#### Nuclear Extracts (High salt)

For nuclear extract preparation, cells were incubated with buffer A (10mM Hepes pH 7.9, 2.5mM MgCl2, 0.25M sucrose, 0.1% NP40 and protease inhibitors) for 10 min on ice, centrifuged at 8000 rpm for 10 min, resuspended in buffer B (25mM Hepes pH 7.9, 1.5mM MgCl2, 700 mM NaCl, 0.1 mM EDTA, 20% glycerol and protease inhibitors), sonicated and centrifuged at 14000 rpm for 15min.

#### Chromatography analyses

For analysis of endogenous protein profiles in HAP1 cells, nuclear extracts (High salt) were first dialyzed against BP100 (50mM Potassium Phosphate pH 6,8, 100mM NaCl, 1mM EDTA, 1mM DTT, protease inhibitors) and clarified by high-speed centrifugation. Samples were then purified by ion exchange chromatography using a HiTrap SP HP 5ml column (GE Healthcare). Elution was performed by step elution with increasing NaCl concentration. 500mM elution was concentrated 5X time on centricon (Millipore, cut-off 10kDa) and analyzed on Superose 6 PC 3.2 increase column (GE Healthcare). The native molecular size markers used for column calibration were thyroglobulin (669 kDa), ferritin (440 kDa) and aldolase (158 kDa).

#### Mass spectrometry analysis

For mass spectrometry analysis of Flag-tagged constructs overexpressed in HeLa cells, 100mg of nuclear extracts were used. Nuclear extracts were first dialyzed in BC250 (50mM Tris pH8.0, 250mM KCl, 1mM EDTA, 10% Glycerol and protease inhibitors). Precipitates were removed by centrifugation and the supernatant was incubated with anti-FLAG M2 affinity gel overnight. The beads were then washed three times with BC250 + 0,05% NP40 and eluted with 0,2mg/ml Flag peptide, precipitated with ice-cold acetone and resuspended in 1X Laemmli Sample Buffer. Of note, for mass spectrometry analysis of Flag-tagged BAP1 overexpressed in MP41 cells, 50 mg of nuclear extracts was used and first purified on an ion exchange chromatography HiTrap Q 1ml column (GE Healthcare).

After IP and elution of enriched proteins, SDS-PAGE (Invitrogen) was used without separation as a cleanup step to remove lipids, metabolites, salts, and denaturing agents from the samples. After colloidal blue staining (LabSafe GEL BlueTM GBiosciences), 4 gel slices were excised and proteins were reduced with 10 mM DTT prior to alkylation with 55 mM iodoacetamide. After washing and shrinking the gel pieces with 100% MeCN, in-gel digestion was performed using trypsin/Lys-C (Promega) overnight in 25 mM NH4HCO3 at 30°C.

Peptides were extracted and analyzed by nanoLC-MS/MS using an RSLCnano system (Ultimate 3000, Thermo Scientific) coupled to an Orbitrap Fusion mass spectrometer (Q-OT-qIT, Thermo Fisher Scientific). Samples were loaded on a C18 precolumn (300 μm inner diameter x 5 mm; Dionex) at 20 μl/min in 2% MeCN, 0.05% TFA. After a desalting for 3 min, the precolumn was switched on the C18 column (75 *μ* m i.d. × 50 cm, packed with C18 PepMap™, 3 *μ* m, 100 Å; LC Packings) equilibrated in solvent A (2% MeCN, 0.1% HCOOH). Bound peptides were eluted using a two-step linear gradient of 147 min (from 1 to 20% (v/v)) of solvent B (100% MeCN, 0.085% HCOOH) and 65 min (from 20 to 40% (v/v)) of solvent B, at a 400 nl/min flow rate and an oven temperature of 40° C. We acquired Survey MS scans in the Orbitrap on the 400–1200 m/z range with the resolution set to a value of 120,000 and a 4 × 10^5^ ion count target. Each scan was recalibrated in real time by co-injecting an internal standard from ambient air into the C-trap. Tandem MS was performed by isolation at 1.6 Th with the quadrupole, HCD fragmentation with normalized collision energy of 35, and rapid scan MS analysis in the ion trap. The MS2 ion count target was set to 10^4^ and the max injection time was 100 ms. Only those precursors with charge state 2–7 were sampled for MS2. The dynamic exclusion duration was set to 60s with a 10ppm tolerance around the selected precursor and its isotopes. The instrument was run in top speed mode with 3 s cycles.

Data were searched against the uniprot-Human database, using Sequest HT from Proteome Discoverer 1.4 (thermo Scientific). Enzyme specificity was set to trypsin and a maximum of two miss cleavages was allowed. Oxidized methionine, N-terminal acetylation and carbamidomethyl cysteine were set as variable modifications. The mass tolerances in MS and MS/MS were set to 10 ppm and 0.6 Da, respectively. The resulting files were further processed using myProMS ^56^. The Sequest HT target and decoy search results were validated at 1% FDR with Percolator.

#### DNA methylation analysis

Genomic DNA was extracted following manufacturer protocol (DNA easy, Qiagen), treated with RNAse A and RNAse T1, purified again and quantified using nanodrop. 1 microgram of DNA was then digested by the degradase plus (Zymoresearch), ethanol precipitated and the supernatant was then evaporated on speedvac. Samples were then reconstituted in 10 μl of solution A’ (2% methanol, 0.1% HCOOH), vortex-mixed, centrifuged and transferred to a HPLC vial for micoLC-MS/MS analysis. Fractions were used directly in solution A’ and analyzed (5 μL) using the RSLCnano system connected to the Orbitrap Fusion mass spectrometer. Sample separation was achieved on a C18 column (4.6 × 100 mm, packed with ZORBAX Eclipse XDB C18, 1.8 μm particles, Agilent Technologies) after 5 min loading in solvent A’, with a linear gradient of 10 min (from 0 to 30% (v/v) of solvent B’ (80% MeCN, 0.1% HCOOH)) at 500 μL/min. Data acquisition was performed in the Orbitrap on the 200–300 m/z range with the resolution set to a value of 240,000 at m/z 200. To determine the intensity of each nucleosides, we extracted from the MS survey of microLC-MS/MS raw files the extracted ion chromatogram (XIC) signal by using the retention time and m/z values of the well characterized synthetic nucleoside ions using the Xcalibur softwares (manually). XIC areas were integrated in Xcalibur under the QualBrowser interface using the ICIS algorithm.

## Acknowledgements

The Labex DEEP, AAP EpiG and Cancer, “Programme Incitatif et Coopératif” (PIC) Uveal Melanoma and the Institut Curie supported work in R.M. laboratory. A.C. and M.W. were recipients of fellowships from the “Association pour la Recherche contre le Cancer (ARC)”. Mass spectrometry analyses were performed by the Institut Curie Protein Mass Spectrometry Laboratory, supported by grants from “Region Ile-de-France” and the FRM. High-throughput sequencing was performed by the NGS platform of the Institut Curie, supported by grants ANR-10-EQPX-03 and ANR10-INBS-09–08 from the Agence Nationale de le Recherche (investissements d’avenir) and by the Canceropôle Ile-de-France. We thank Tatiana Popova for help with the RNA-seq and, members of the Margueron lab, M-H Stern and P. Gilardi for comments on the manuscript.

## Bibliography

1. Jensen, D.E. et al. BAP1: a novel ubiquitin hydrolase which binds to the BRCA1 RING finger and enhances BRCA1-mediated cell growth suppression. Oncogene 16, 1097–112 (1998).

2. Carbone, M. et al. BAP1 and cancer. Nat Rev Cancer 13, 153–9 (2013).

3. Yu, H. et al. The ubiquitin carboxyl hydrolase BAP1 forms a ternary complex with YY1 and HCF-1 and is a critical regulator of gene expression. Mol Cell Biol 30, 5071–85 (2010).

4. Dey, A. et al. Loss of the tumor suppressor BAP1 causes myeloid transformation. Science 337, 1541–6 (2012).

5. Machida, Y.J., Machida, Y., Vashisht, A.A., Wohlschlegel, J.A. & Dutta, A. The deubiquitinating enzyme BAP1 regulates cell growth via interaction with HCF-1. J Biol Chem 284, 34179–88 (2009).

6. Nishikawa, H. et al. BRCA1-associated protein 1 interferes with BRCA1/BARD1 RING heterodimer activity. Cancer Res 69, 111–9 (2009).

7. Ji, Z. et al. The forkhead transcription factor FOXK2 acts as a chromatin targeting factor for the BAP1-containing histone deubiquitinase complex. Nucleic Acids Res 42, 6232–42 (2014).

8. Baymaz, H.I. et al. MBD5 and MBD6 interact with the human PR-DUB complex through their methyl-CpG-binding domain. Proteomics 14, 2179–89 (2014).

9. Hauri, S. et al. A High-Density Map for Navigating the Human Polycomb Complexome. Cell Rep 17, 583–595 (2016).

10. Zarrizi, R., Menard, J.A., Belting, M. & Massoumi, R. Deubiquitination of gamma-tubulin by BAP1 prevents chromosome instability in breast cancer cells. Cancer Res 74, 6499–508 (2014).

11. Lee, H.S., Lee, S.A., Hur, S.K., Seo, J.W. & Kwon, J. Stabilization and targeting of INO80 to replication forks by BAP1 during normal DNA synthesis. Nat Commun 5, 5128 (2014).

12. Matatall, K.A. et al. BAP1 deficiency causes loss of melanocytic cell identity in uveal melanoma. BMC Cancer 13, 371 (2013).

13. Scheuermann, J.C. et al. Histone H2A deubiquitinase activity of the Polycomb repressive complex PR-DUB. Nature 465, 243–7 (2010).

14. Gaytan de Ayala Alonso, A. et al. A genetic screen identifies novel polycomb group genes in Drosophila. Genetics 176, 2099–108 (2007).

15. Simon, J.A. & Kingston, R.E. Mechanisms of polycomb gene silencing: knowns and unknowns. Nat Rev Mol Cell Biol 10, 697–708 (2009).

16. Margueron, R. & Reinberg, D. The Polycomb complex PRC2 and its mark in life. Nature 469, 343–9 (2011).

17. Wang, H. et al. Role of histone H2A ubiquitination in Polycomb silencing. Nature 431, 873–8 (2004).

18. Pengelly, A.R., Kalb, R., Finkl, K. & Muller, J. Transcriptional repression by PRC1 in the absence of H2A monoubiquitylation. Genes Dev 29, 1487–92 (2015).

19. Illingworth, R.S. et al. The E3 ubiquitin ligase activity of RING1B is not essential for early mouse development. Genes Dev 29, 1897–902 (2015).

20. Blackledge, N.P. et al. Variant PRC1 complex-dependent H2A ubiquitylation drives PRC2 recruitment and polycomb domain formation. Cell 157, 1445–59 (2014).

21. Cooper, S. et al. Targeting polycomb to pericentric heterochromatin in embryonic stem cells reveals a role for H2AK119u1 in PRC2 recruitment. Cell Rep 7, 1456–1470 (2014).

22. Scheuermann, J.C., Gutierrez, L. & Muller, J. Histone H2A monoubiquitination and Polycomb repression: the missing pieces of the puzzle. Fly (Austin) 6, 162–8 (2012).

23. Sahtoe, D.D., van Dijk, W.J., Ekkebus, R., Ovaa, H. & Sixma, T.K. BAP1/ASXL1 recruitment and activation for H2A deubiquitination. Nat Commun 7, 10292 (2016).

24. Abdel-Wahab, O. et al. ASXL1 mutations promote myeloid transformation through loss of PRC2-mediated gene repression. Cancer Cell 22, 180–93 (2012).

25. Abdel-Wahab, O. et al. Deletion of Asxl1 results in myelodysplasia and severe developmental defects in vivo. J Exp Med 210, 2641–59 (2013).

26. Inoue, D. et al. Myelodysplastic syndromes are induced by histone methylation-altering ASXL1 mutations. J Clin Invest 123, 4627–40 (2013).

27. Li, T., Hodgson, J.W., Petruk, S., Mazo, A. & Brock, H.W. Additional sex combs interacts with enhancer of zeste and trithorax and modulates levels of trimethylation on histone H3K4 and H3K27 during transcription of hsp70. Epigenetics Chromatin 10, 43 (2017).

28. Valletta, S. et al. ASXL1 mutation correction by CRISPR/Cas9 restores gene function in leukemia cells and increases survival in mouse xenografts. Oncotarget 6, 44061–71 (2015).

29. LaFave, L.M. et al. Loss of BAP1 function leads to EZH2-dependent transformation. Nat Med 21, 1344–9 (2015).

30. Wu, X. et al. Tumor suppressor ASXL1 is essential for the activation of INK4B expression in response to oncogene activity and anti-proliferative signals. Cell Res 25, 1205–18 (2015).

31. Oak, J.S. & Ohgami, R.S. Focusing on frequent ASXL1 mutations in myeloid neoplasms, and considering rarer ASXL2 and ASXL3 mutations. Curr Med Res Opin 33, 781–782 (2017).

32. Carette, J.E. et al. Global gene disruption in human cells to assign genes to phenotypes by deep sequencing. Nat Biotechnol 29, 542–6 (2011).

33. Wassef, M. et al. Versatile and precise gene-targeting strategies for functional studies in mammalian cell lines. Methods 121–122, 45–54 (2017).

34. Daou, S. et al. The BAP1/ASXL2 Histone H2A Deubiquitinase Complex Regulates Cell Proliferation and Is Disrupted in Cancer. J Biol Chem 290, 28643–63 (2015).

35. Almeida, M. et al. PCGF3/5-PRC1 initiates Polycomb recruitment in X chromosome inactivation. Science 356, 1081–1084 (2017).

36. Riising, E.M. et al. Gene silencing triggers polycomb repressive complex 2 recruitment to CpG islands genome wide. Mol Cell 55, 347–60 (2014).

37. Schoumacher, M. et al. Uveal melanoma cells are resistant to EZH2 inhibition regardless of BAP1 status. Nature Medicine 22(2016).

38. Lee, S.W. et al. ASXL1 represses retinoic acid receptor-mediated transcription through associating with HP1 and LSD1. J Biol Chem 285, 18–29 (2010).

39. Park, U.H., Yoon, S.K., Park, T., Kim, E.J. & Um, S.J. Additional sex comb-like (ASXL) proteins 1 and 2 play opposite roles in adipogenesis via reciprocal regulation of peroxisome proliferator-activated receptor {gamma}. J Biol Chem 286, 1354–63 (2011).

40. Dubey, R. et al. Chromatin-Remodeling Complex SWI/SNF Controls Multidrug Resistance by Transcriptionally Regulating the Drug Efflux Pump ABCB1. Cancer Res 76, 5810–5821 (2016).

41. Balasubramani, A. et al. Cancer-associated ASXL1 mutations may act as gain-of-function mutations of the ASXL1-BAP1 complex. Nat Commun 6, 7307 (2015).

42. Klymenko, T. & Muller, J. The histone methyltransferases Trithorax and Ash1 prevent transcriptional silencing by Polycomb group proteins. EMBO Rep 5, 373–7 (2004).

43. Sinclair, D.A., Campbell, R.B., Nicholls, F., Slade, E. & Brock, H.W. Genetic analysis of the additional sex combs locus of Drosophila melanogaster. Genetics 130, 817–25 (1992).

44. Fisher, C.L. et al. Additional sex combs-like 1 belongs to the enhancer of trithorax and polycomb group and genetically interacts with Cbx2 in mice. Dev Biol 337, 9–15 (2010).

45. Gutierrez, L. et al. The role of the histone H2A ubiquitinase Sce in Polycomb repression. Development 139, 117–27 (2012).

46. Sanchez, R. & Zhou, M.M. The PHD finger: a versatile epigenome reader. Trends Biochem Sci 36, 364–72 (2011).

47. Nakatani, Y. & Ogryzko, V. Immunoaffinity purification of mammalian protein complexes. Rna Polymerases and Associated Factors, Pt C 370, 430–444 (2003).

48. Hebert, L. et al. Modulating BAP1 expression affects ROS homeostasis, cell motility and mitochondrial function. Oncotarget 8, 72513–72527 (2017).

49. martin, M. Cutadapt removes adapter sequences from high-throughput sequencing reads. EMBnet.journal 17, 10–12 (2011).

50. Langmead, B., Trapnell, C., Pop, M. & Salzberg, S.L. Ultrafast and memory-efficient alignment of short DNA sequences to the human genome. Genome Biol 10, R25 (2009).

51. Dobin, A. et al. STAR: ultrafast universal RNA-seq aligner. Bioinformatics 29, 15–21 (2013).

52. Liao, Y., Smyth, G.K. & Shi, W. featureCounts: an efficient general purpose program for assigning sequence reads to genomic features. Bioinformatics 30, 923–30 (2014).

53. Li, B. & Dewey, C.N. RSEM: accurate transcript quantification from RNA-Seq data with or without a reference genome. BMC Bioinformatics 12, 323 (2011).

54. Wang, L., Wang, S. & Li, W. RSeQC: quality control of RNA-seq experiments. Bioinformatics 28, 2184–5 (2012).

55. Margueron, R. et al. Ezh1 and Ezh2 maintain repressive chromatin through different mechanisms. Mol Cell 32, 503–18 (2008).

56. Poullet, P., Carpentier, S. & Barillot, E. myProMS, a web server for management and validation of mass spectrometry-based proteomic data. Proteomics 7, 2553–6 (2007).

